# Type-I IFNs in EAE drive synaptic transcriptional responses in neurons and astrocytes in the brain

**DOI:** 10.64898/2026.09.23.753833

**Authors:** Shane M. O’Neil, Hyunjung Min, Li Xu, Anthony J. Filiano

## Abstract

Neuronal pathology plays a prominent role in the cognitive and neuropathic symptoms of multiple sclerosis; however, little is known about these pathological signals to neurons in the brain and their consequences on cognition. Here, using a transgenic reporter of IFN signaling with spatial RNA-sequencing, we determined myelin-specific autoimmune T cells enter the brain parenchyma through the interface between the fimbria and third ventricle. Here, astrocytes and microglia express high levels of MHC-II-associated transcripts. Using this reporter, we also showed microglia within the brain are not direct responders of IFNs, but express transcripts associated with synaptic pruning, potentially contributing to pathological remodeling of neuronal circuits. Moreover, both type-I and type-II IFNs are required for T cells to infiltrate the brain and initiate an IFN response associated with synaptic and metabolic programs in neurons and astrocytes in the cortex and hippocampus. In summary, this study elucidates multiple steps of the pathogenesis of EAE pathology within the brain independent of the spinal cord.

IFNs activate astrocytes and neurons in EAE brain

**Summary:** Autoreactive T cells in EAE activate IFN-responsive pathways in neurons and astrocytes in the brain. T cells enter the brain through the fimbria and drive transcriptional signatures of synaptic dysfunction in neurons and synaptic pruning in microglia.

## Introduction

Multiple sclerosis (MS), a chronic, demyelinating, autoimmune disorder characterized by inflammation, gliosis, and neurodegeneration, affects nearly one million people in the United States (Wallin et al., 2019). Experimental autoimmune encephalomyelitis (EAE), the most common animal model of MS, recapitulates the disease using encephalitogenic T cells primed against the myelin oligodendrocyte glycoprotein (MOG)_35-55_ peptide (Bjelobaba et al., 2018; Schwentker and Rivers 1934; Wolf et al., 1947). Ascending paralysis begins in mice immunized for EAE within two weeks as inflammation and leukocyte infiltration progress up the spinal cord toward the brain. Studies have demonstrated that endothelial barrier breakdown occurs in the presymptomatic stage of EAE but is limited to the meninges (Politi et al., 2007). Others, however, have shown sparse infiltration of T cells into the striatum closer, but still prior, to the onset of symptoms (Centonze et al., 2009). As the cognitive symptoms, such as anxiety, depression, and neuropathic pain, often go neglected in the clinic, the neuropathology within the cortex of MS patients and EAE mice, whether independent of or a part of demyelination, deserves more thorough investigation (Margoni et al., 2023).

Interferon (IFN) signaling, a key component of an organism’s response to viral infection, is highly implicated in the pathophysiology of MS and other autoimmune disorders (Khan et al., 2024; Wang et al., 2024). Autoreactive, IFN-γ-producing T cells are commonly found within white matter active lesions in MS and EAE (Aboelnour et al., 2026). Early in disease progression, cortical inflammation is associated with cognitive impairment and delirium (Zupo et al., 2026). This cortical inflammation is independent of infiltrating leukocytes and is suspected to be driven by glia responding to cytokines released by inflammatory cells in the overlying meninges (Florescu et al., 2025; Naouar et al., 2026); however, the mechanism and signaling events of this process are unclear.

In this study, we identified spatially distinct IFN-responsive astrocytes and excitatory neurons in the somatosensory (SS) cortex and CA1 region of the hippocampus at the onset of EAE symptoms. Additionally, we found microglia expressing transcripts associated with synaptic pruning within these tissues. Using passive EAE with 2D2 T cells, we identified the medial edge of the fimbria as a key location for central nervous system (CNS) antigen-specific T cells to infiltrate the brain parenchyma. Moreover, the advancement of transgenic and spatial transcriptomic techniques has enabled unbiased investigation into sparse transcriptional effects scattered across a tissue sample (Bonev et al., 2024). Thus, we investigated the transcriptional and cellular profiles of various brain regions. Notably, we isolated IFN-stimulated cell populations *in silico* for comparison by RNA-sequencing, whereby we identified distinct transcriptional phenotypes of various IFN-stimulated cell populations in the EAE brain.

## Results

### Astrocytes and neurons within the brain react to IFN signaling at the onset of EAE symptoms

To observe IFN signaling within the CNS, we generated Mx-Cre^+^::Ai9^+/-^ reporter mice (MX1^tdT^) in which red-fluorescent tdTomato (tdT) is expressed under the *Mx1* promoter (**Fig. 1A**). *Mx1* is expressed downstream of type-I, -II, and -III IFN signaling; therefore, cells that had been stimulated by IFNs were tdT^+^. We then induced active EAE in these mice and observed their phenotype over 14 days post immunization (DPI). Mice treated with CFA/MOG_35-55_ emulsion (EAE group) lost more weight than controls treated with pertussis toxin (PTX) alone (**Fig. 1B**). Moreover, all EAE mice developed clinical symptoms at 14 DPI (**Fig. 1C**). Brain tissue was collected at the onset of symptoms (or 14 DPI for PTX-treated control mice). At this time, EAE mice had a higher number of tdT^+^ cells in the brain, and these cells were primarily SOX9^+^ astrocytes and NeuN^+^ pyramidal cells within the SS cortex and CA1 region of the hippocampus (**Fig. 1D-G**). Moreover, MX1-tdT expression was higher in the spinal cord of EAE mice and increased posteriorly from cervical to lumbar segments (**Fig.S1A&B**). In contrast to the brain, the MX1-tdT expression in the spinal cord was concentrated in GFAP^+^ astrocytes and IBA1^+^ microglia. Upon closer interrogation, we discovered MX1-tdT fluorescence within microglia in the brain, but this fluorescence was restricted to CD68^+^ phagolysosomes, whereas MX1-tdT fluorescence was observed throughout entire spinal cord microglial cells (**Fig. 1H**). Taken together, these data indicate neurons and astrocytes within the brain respond directly to IFN signaling, while astrocytes and microglia respond to IFN signaling in the spinal cord.

**Figure 1.**
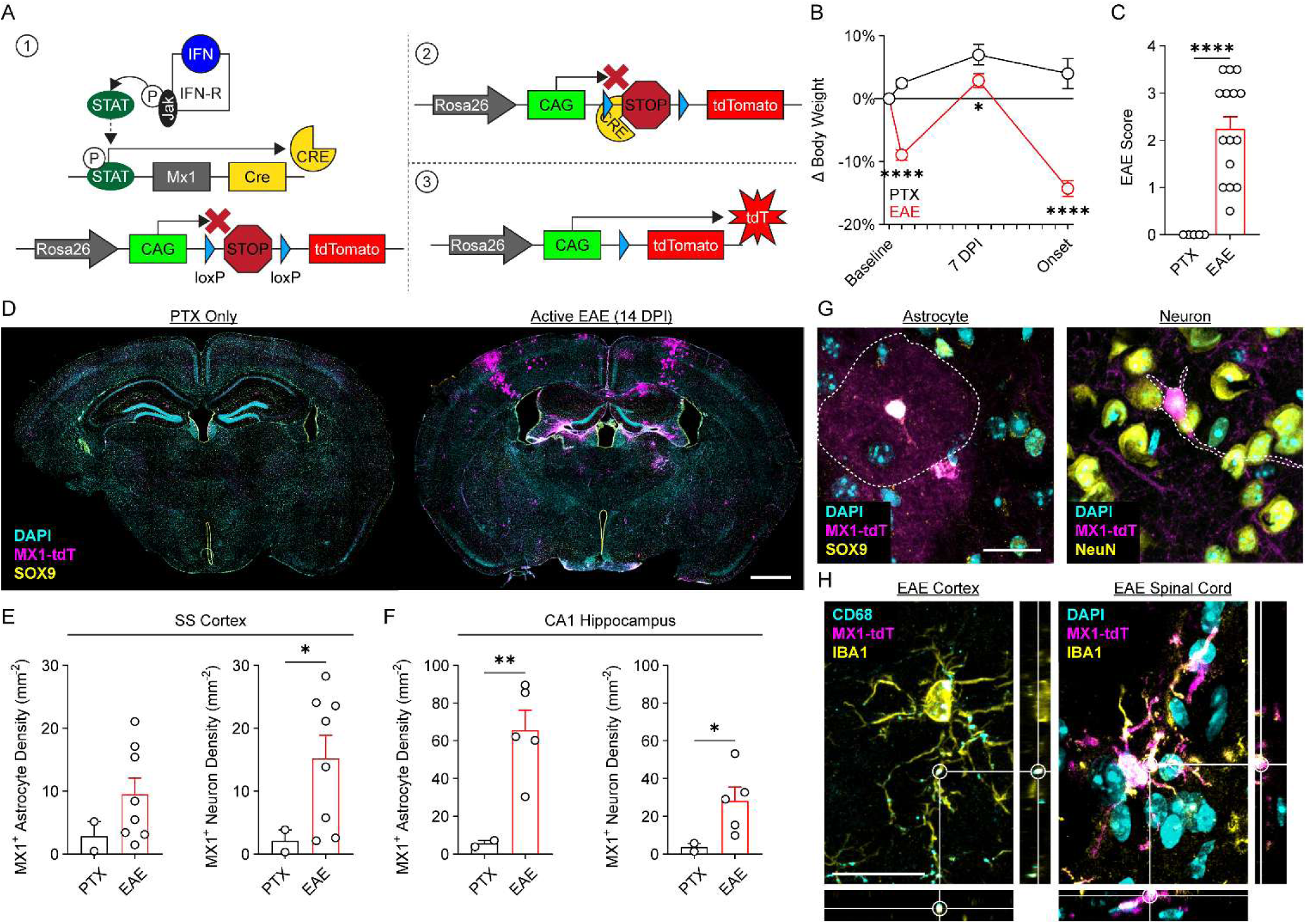
Astrocytes and neurons within the brain react to IFN signaling at the onset of EAE symptoms. (A) Schematic of transgenic tdTomato reporter mechanism in MX1^tdT^ mice. Scale bar = 1 mm. **(B)** Body weight of mice treated with either PTX alone or PTX and MOG35-55/CFA emulsion (EAE). Time × Treatment Interaction: F(3, 26) = 31.65, *P* < 0.0001. **(C)** EAE symptom scores for mice at 14 DPI (symptom onset). **(D)** Representative coronal brain sections from mice at 14 DPI. **(E&F)** Quantification of MX1-tdT^+^ astrocyte and neuron density in the somatosensory (SS) cortex and CA1 region of the hippocampus at 14 DPI. **(G)** Representative images of an individual MX1-tdT^+^ astrocyte (left) and neuron (right). Scale bar = 20 µm. **(H)** Representative orthogonal projections of MX1-tdT fluorescence in the EAE cortex (left) and spinal cord (right). Scale bar = 20 µm. Data shown are mean ± SEM. \**P* < 0.05; \*\**P* < 0.01; \*\*\*\**P* < 0.0001.

### MOG_35-55_-restricted T_h_1 cells traffic along dural sinuses early in passive EAE

To avoid inflammatory effects of EAE priming and focus on the role of IFN-producing T cells, we next induced passive EAE in these MX1^tdT^ reporter mice using 2D2 T helper 1 (T_h_1) cells (**Fig. 2A**). A T_h_1 phenotype was confirmed by expression of CD4, IFN-γ, and T-bet using flow cytometry (**Fig. 2B**). Using CD45.1^+^ 2D2 T cells for identification by flow cytometry, relevant tissues were isolated from recipient mice and analyzed using flow cytometry (**Fig.S2**). At this time, 25% of lymphoid cells (CD45^+^/CD11b^-^) in the dura mater were donor 2D2 cells, which was a greater proportion compared to all other tissues measured (**Fig. 2C**). Based on these results, we repeated this experiment using GFP^+^ 2D2 T_h_1 cells and analyzed the dura mater using immunohistochemistry. Here, we observed an increase in GFP^+^ 2D2 T cells along the superior sagittal and transverse sinuses from 3 to 7 DPI (**Fig. 2D&E**). Finally, we confirmed induction of MX1-tdT expression in passive EAE in the spinal cord (**Fig.S1C&D**) and astrocytes and neurons within the SS cortex and CA1 hippocampus at the onset of EAE symptoms, as was observed with active EAE (**Fig.2F-H**). Taken together, these data show CNS antigen-specific T_h_1 cells traffic along the dural sinuses prior to the onset of EAE symptoms.

**Figure 2.**
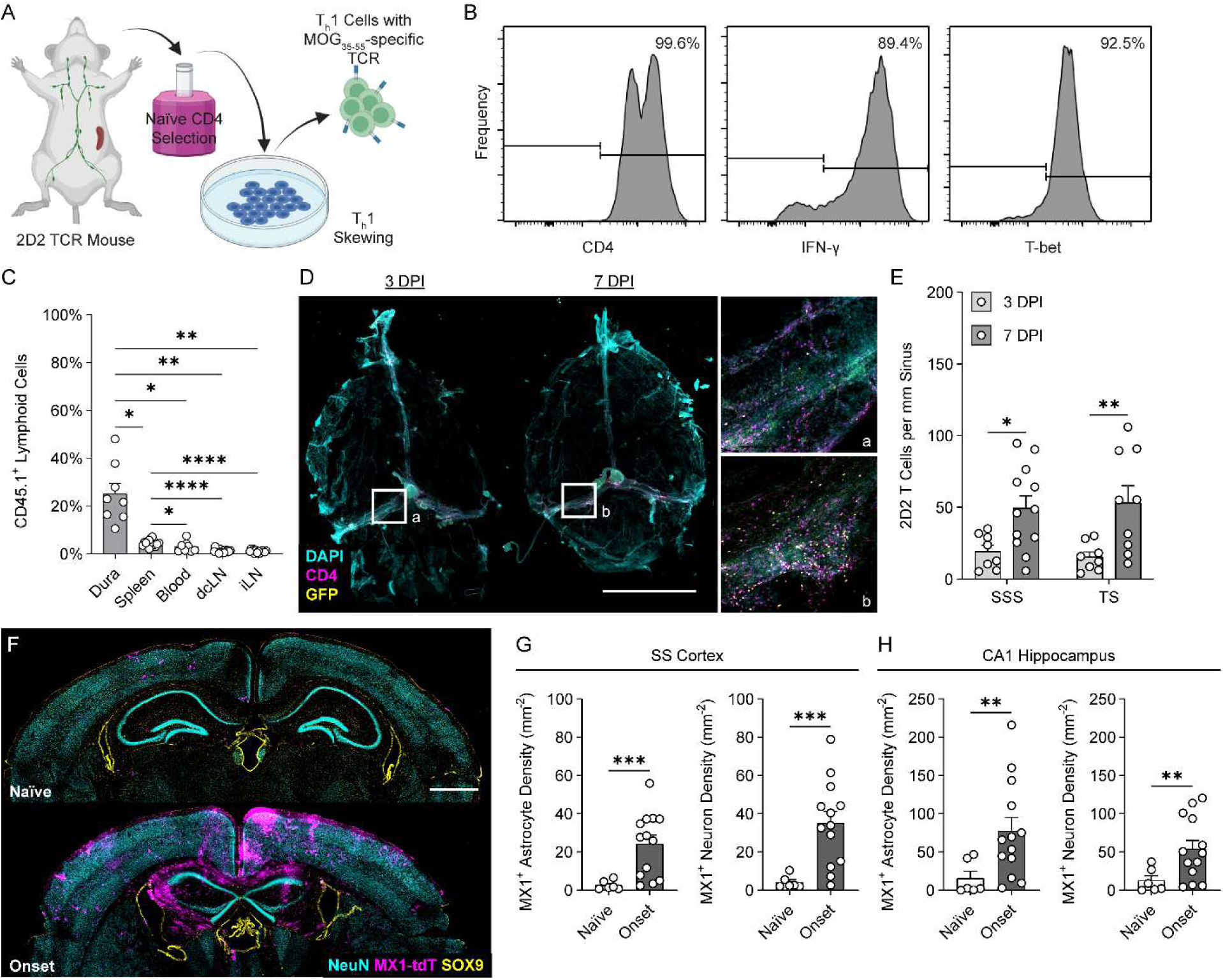
MOG35-55-restricted Th1 cells traffic along dural sinuses early in passive EAE. **(A)** Schematic of 2D2 T cell isolation, culture, and adoptive transfer for passive EAE. **(B)** Representative flow cytometry histograms confirming Th1 phenotype. **(C)** Quantification of labeled 2D2 cells detected in each tissue by flow cytometry compared to the total number of lymphocytes in each tissue at 5 DPI. Main Effect of Treatment: F(1, 11) = 44.59, *P* < 0.0001. **(D)** Representative images of dura mater whole-mounts at 3 and 7 DPI. Scale bar = 5 mm. **(E)** Quantification of GFP^+^ 2D2 T cells located along the superior sagittal sinus (SSS) and transverse sinuses (TS) in the dura mater. Main Effect of Time: F(1, 33) = 16.42, *P* = 0.0003. **(F)** Representative coronal brain sections from mice at 12 DPI (symptom onset) after induction of passive EAE. Scale bar = 1 mm. **(G&H)** Quantification of MX1-tdT^+^ astrocyte and neuron density in the SS cortex and CA1 region of the hippocampus at symptom onset. Data shown are mean ± SEM. \**P* < 0.05; \*\**P* < 0.01; \*\*\**P* < 0.001; \*\*\*\**P* < 0.0001.

### IFN-dependent dysfunction of synaptic genes in excitatory neurons in the brain at EAE symptom onset is correlated with synaptic pruning signatures in microglia

We next used high-resolution spatial RNA transcriptomic analysis to determine spatially distinct effects within key regions of the brain as a result of passive EAE. Focusing on the SS cortex, a region in which a large population of MX1-tdT^+^ astrocytes and neurons were observed, we identified seven key cell types between naïve and EAE (symptom onset) samples (**Fig. 3A-C**). To focus on IFN-induced effects within the SS cortex, EAE-dependent differentially expressed genes (DEGs) within each cell type were filtered to remove effects shared by the thalamus, a region lacking MX1-tdT expression throughout experiments (**Fig. 3D**). This filter yielded lists of DEGs with a large focus on excitatory neurons and oligodendrocytes (**Fig. 3E**). Within excitatory neurons, these DEGs revealed an upregulation of processes including long-term synaptic depression (GO:0060292), synaptic vesicle maturation (GO:0016188), postsynaptic density organization (GO:0097106), and regulation of synapse organization (GO:0050807) alongside a downregulation of clathrin coat disassembly (GO:0072318), regulation of calcium ion transmembrane transport (GO:1903169), regulation of membrane potential (GO:0042391), and modulation of chemical synapse transmission (GO:0050804), indicating an overall dysregulation of transsynaptic activity in these neurons (**Fig. 3F&G**).

**Figure 3.**
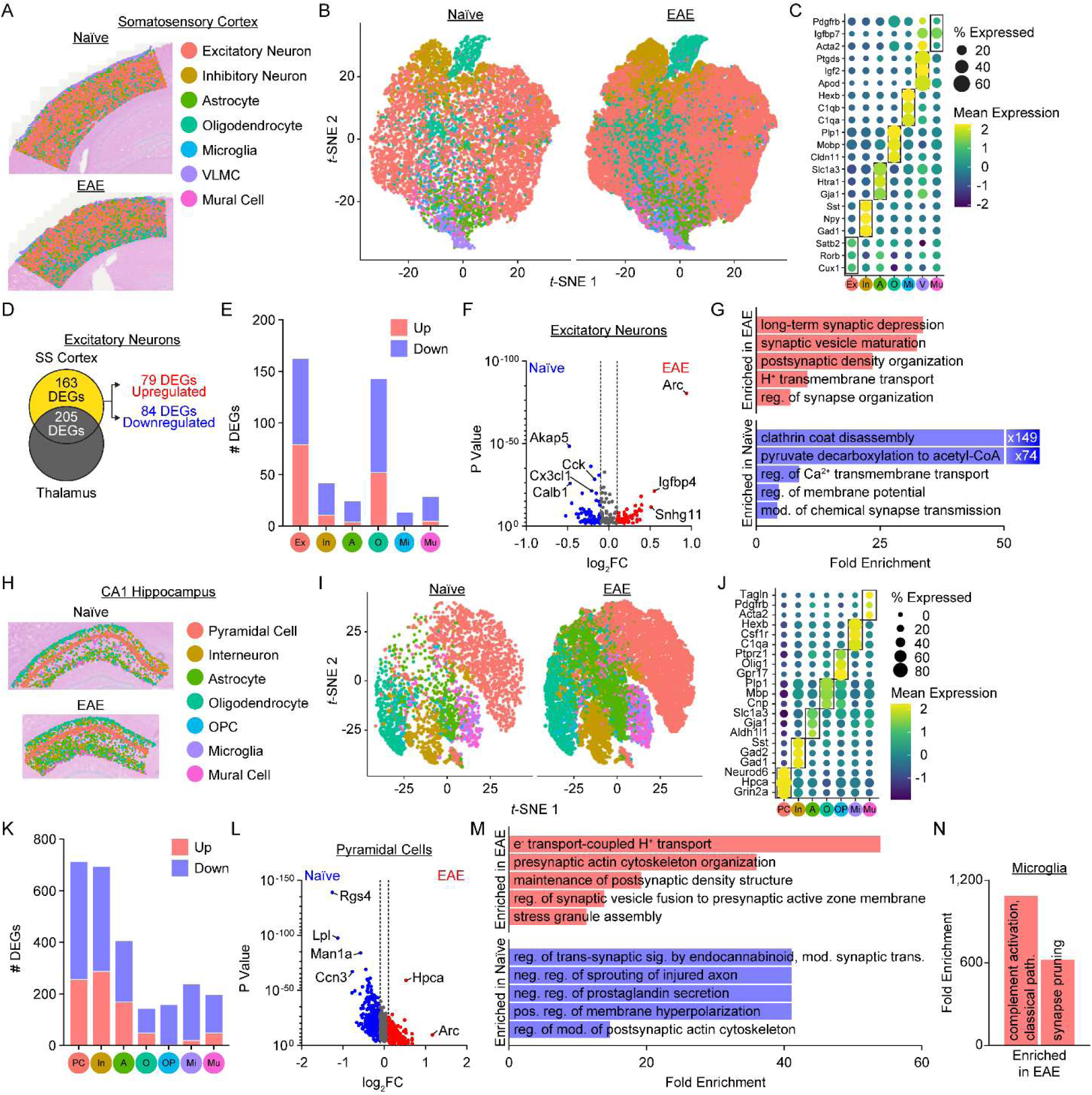
IFN-dependent dysfunction of synaptic genes in excitatory neurons in the brain at EAE symptom onset is correlated with synaptic pruning signatures in microglia. **(A)** Representative spatial plots from Visium HD spatial RNA sequencing of the SS cortex. **(B)** Cell types and clustering of cells shown by *t*-SNE plot. **(C)** Representative signature gene expression used to identify cell types. **(D)** Schematic of DEG filtering by removing DEGs shared by the thalamus, a region with minimal MX1-tdT expression. **(E)** Number of filtered up- and down-regulated DEGs with EAE by cell type. **(F)** Volcano plot of filtered DEGs in excitatory neurons. **(G)** Significant enriched GO Biological Processes for filtered excitatory neuron DEGs. **(H)** Representative spatial plots of the CA1 region of the hippocampus. **(I)** Cell types and clustering of cells shown by *t*-SNE plot. **(J)** Representative signature gene expression used to identify cell types. **(K)** Number of filtered up- and down-regulated DEGs with EAE by cell type. **(L)** Volcano plot of filtered DEGs in pyramidal cells. **(M&N)** Significant enriched GO Biological Processes for filtered pyramidal cell and microglia DEGs.

Shifting focus to the CA1 region of the hippocampus, another region in which a large population of MX1-tdT^+^ astrocytes and neurons were observed, we again identified seven key cell types between naïve and EAE samples (**Fig. 3H-J**). Here, filtered DEGs showed a focus on neurons (excitatory pyramidal cells and inhibitory interneurons) and astrocytes (**Fig. 3K**). Notably, the synaptic gene *Arc* had the highest EAE fold-change in excitatory neurons in both the cortex and hippocampus (**Fig. 3F&L**). CA1 pyramidal neurons upregulated processes including presynaptic actin cytoskeleton organization (GO:0099140), maintenance of postsynaptic density structure (GO:0099562), and regulation of synaptic vesicle fusion to presynaptic active zone membrane (GO:0031630) alongside a downregulation of the regulation of trans-synaptic signaling by endocannabinoid, modulating synaptic transmission (GO:0150036), positive regulation of membrane hyperpolarization (GO:1902632), and regulation of modification of postsynaptic actin cytoskeleton (GO:1905274), again indicating a dysregulation of synaptic function (**Fig. 3L&M**). Interestingly, microglia in this region exhibited a highly specific increase in classical complement-mediated synaptic pruning (**Fig. 3N**). This finding is also consistent with the presence of *Mx1* promoter driven tdT within microglial phagolysosomes (**Fig. 1H**). These data support the synaptic pruning of IFN-stimulated neurons by microglia, leading to synaptic dysfunction of cortical and hippocampal excitatory neurons as a result of EAE.

### IFN-stimulated neurons express transcripts associated with synaptic dysfunction and neurodegeneration

To determine the localized effects of IFN-signaling in the SS cortex and CA1 hippocampus, cell segmentation was performed using fluorescent labeling of consecutive brain sections (**Fig. 4A&E**). Within these tdT^+^ segmentations in the SS cortex, excitatory neurons exhibited an upregulation of transcripts associated with the cellular response to both IFN-β (GO:0035458) and type-II IFN (GO:0071346), validating that MX1-tdT expression in our model is indeed associated with IFN signaling to these cells (**Fig. 4B&C**). Additionally, these neurons upregulated electron transport-coupled proton transport (GO:0015990), nitric oxide-cGMP-mediated signaling (GO:0038060), and calcineurin-NFAT signaling cascade (GO:0033173) and downregulated calcium-regulated lysosome exocytosis (GO:1990927), clathrin coat disassembly (GO:0072318), regulation of clathrin-dependent endocytosis (GO:2000369), calcium ion-regulated exocytosis of neurotransmitter (GO:0048791), and synaptic vesicle endocytosis (GO:0048488). Moreover, oligodendrocytes in these regions, though not tdT^+^, also upregulate the cellular responses to IFN-β and -γ, as well as positive regulation of receptor signaling pathway via JAK-STAT (GO:0046427), the key mediator of IFN receptor signaling within a cell (**Fig. 4D**). In the hippocampus, CA1 pyramidal cells upregulate the positive regulation of neurofibrillary tangle assembly (GO:1902998), negative regulation of amyloid-β formation (GO:1902430), regulation of amyloid-β clearance (GO:1900221), and maintenance of postsynaptic specialization structure (GO:0098880), indicating a phenotype similar to the neurodegeneration seen in disorders such as Alzheimer’s disease (**Fig. 4F&G**). These pyramidal cells also downregulated dendrite extension (GO:0022604), both cholinergic (GO:0032222) and GABAergic (GO:0051932) synaptic transmission, and the ionotropic glutamate receptor signaling pathway (GO:0035235). In CA1 astrocytes, which also expressed MX1-tdT, several mitochondria-associated biological processes were upregulated (*e.g.*, proton motive force-driven mitochondrial ATP synthesis (GO:0042776), mitochondrial electron transport, NADH to ubiquinone (GO:00019646), and regulation of calcium import into the mitochondria (GO:0036444)), representing an increased metabolic demand on IFN-stimulated astrocytes in the CA1 (**Fig. 4H**).

**Figure 4.**
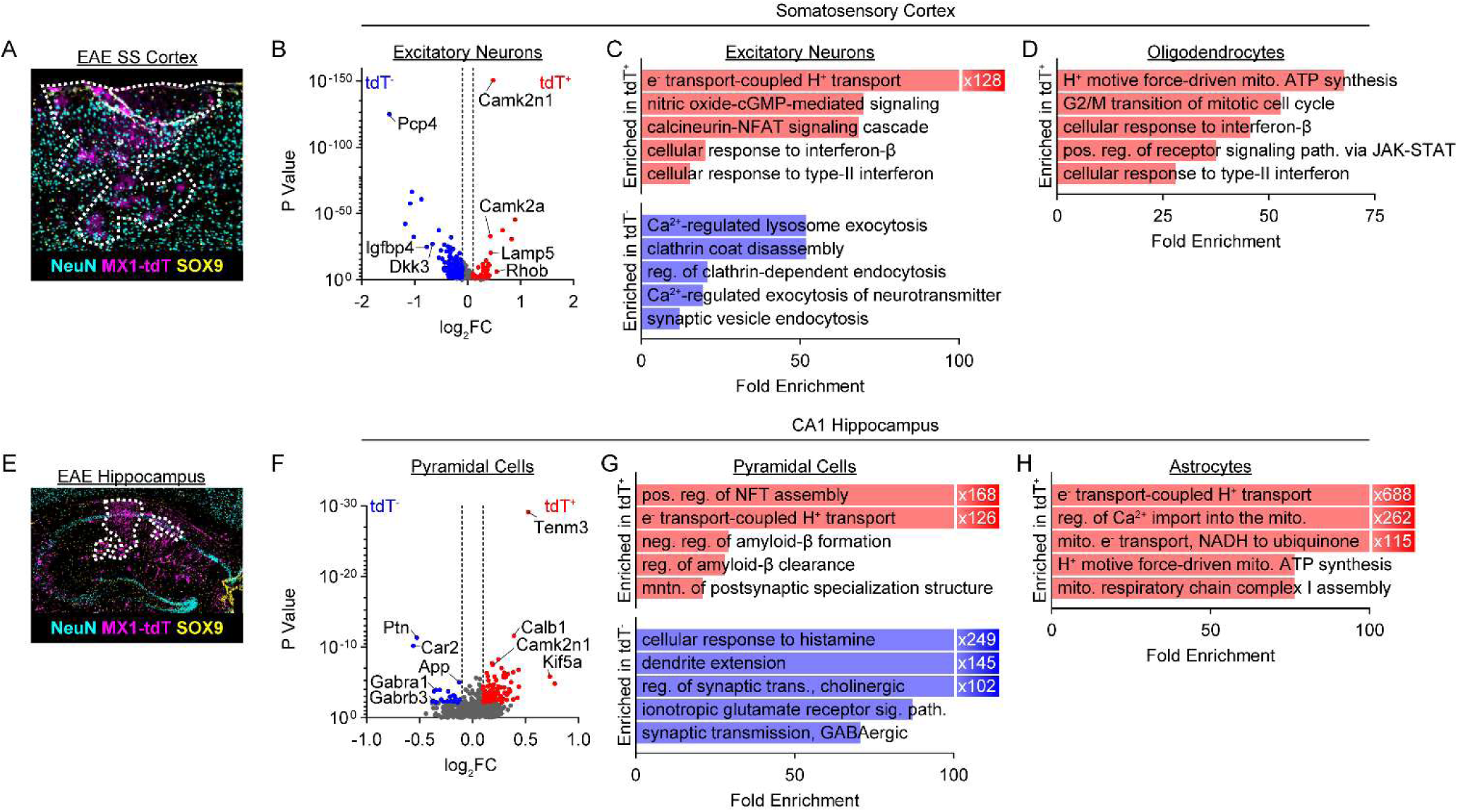
IFN-stimulated neurons express transcripts associated with synaptic dysfunction and neurodegeneration. **(A)** Representative image of segmentation drawn on a consecutive section to identify MX1-tdT-enriched areas in the SS cortex. **(B)** Volcano plot of DEGs in SS cortex MX1-tdT^+^ excitatory neurons. **(C&D)** Significant enriched GO Biological Processes for excitatory neuron and oligodendrocyte DEGs. **(A)** Representative image of segmentation drawn on a consecutive section to identify MX1-tdT-enriched areas in the CA1 region of the hippocampus. **(F)** Volcano plot of DEGs in CA1 MX1-tdT^+^ pyramidal cells. **(G&H)** Significant enriched GO Biological Processes for pyramidal cells and astrocyte DEGs.

### CD4^+^ T cells infiltrate the fimbria at the onset of EAE symptoms

To determine the role of T cells in astrocytic and neuronal IFN-signaling in the brain, we quantified the infiltration of CD4^+^ T_h_ cells into key white and gray matter regions of the brain in EAE (**Fig.5A**). Within the optic nerve and fimbria, and to a lesser degree the corpus callosum and anterior commissure, EAE induced an influx of CD4^+^ T cells at the onset of symptoms (**Fig. 5B**). In addition to these white matter tracts, CD4^+^ T cells also infiltrated the septum and hypothalamus, but not the SS cortex. MX1-tdT expression could also be observed at the onset of EAE symptoms in the optic nerve, fimbria, corpus callosum, and SS cortex (**Fig. 5C**).

**Figure 5.**
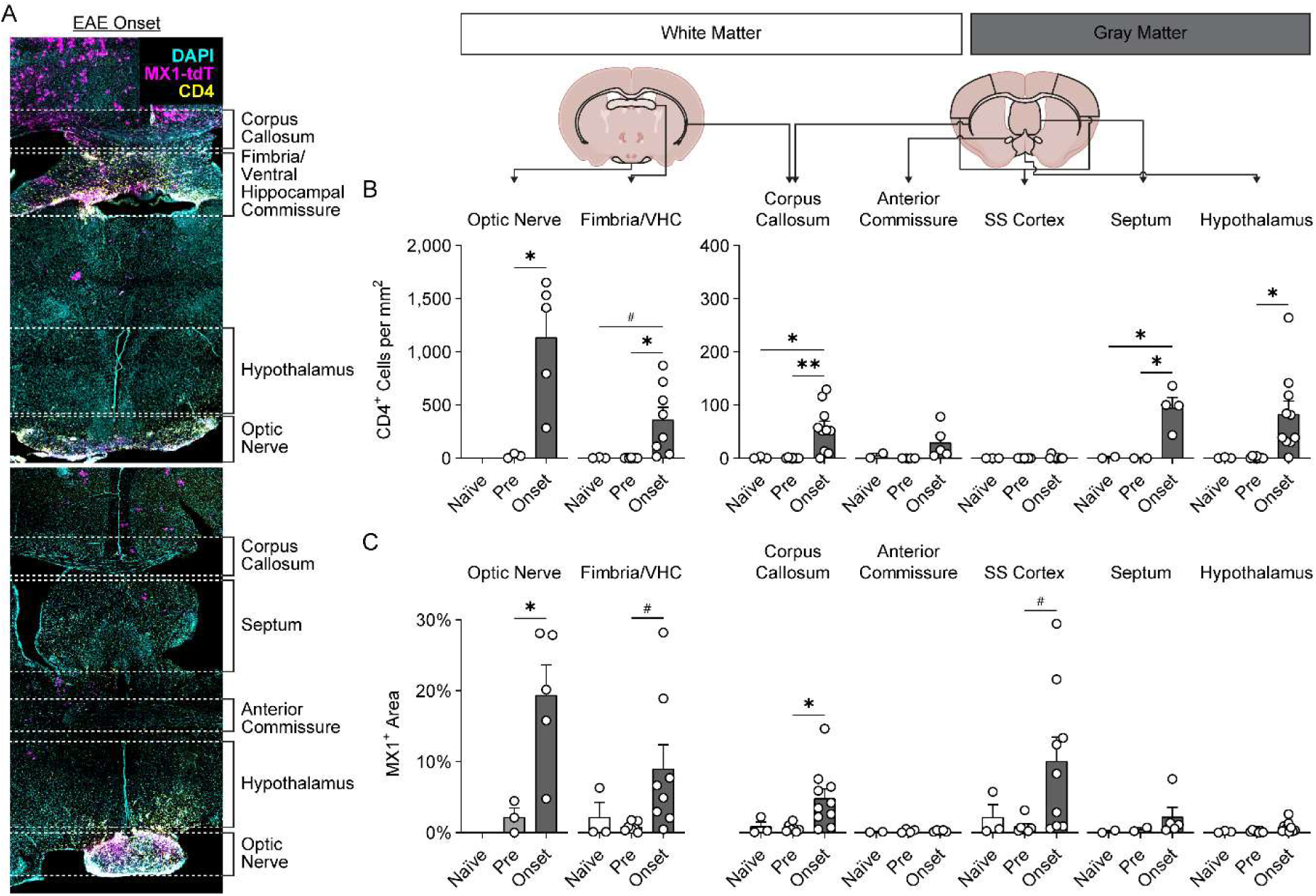
CD4^+^ T cells infiltrate the fimbria at the onset of EAE symptoms. **(A)** Representative coronal brain sections from mice taken at 12 DPI (symptom onset) after induction of passive EAE. **(B&C)** Quantification of CD4^+^ T cell infiltration and MX1-tdT expression in various white matter and gray matter regions at 5 (pre-symptomatic) or 12 DPI. Data shown are mean ± SEM. \**P* < 0.05; \*\**P* < 0.01; ^#^*P* < 0.10.

### IFN-γ is necessary for EAE spinal cord pathology, but type-I IFN signaling is necessary for astrocyte and neuron pathology in the brain

To determine which IFN types were responsible for the MX1-tdT expression in the CNS, we generated *Ifnar1^KO^* and *Ifngr1^KO^* MX1^tdT^ reporter mice and induced passive EAE as before. While we observed MX1-tdT signal within the astrocytes and neurons of the cortex and hippocampus of control mice at the onset of EAE, neither *Ifnar1^KO^*nor *Ifngr1^KO^* mice showed an induction of IFN signaling at the same time point (**Fig. 6A&B**). Moreover, while *Ifngr1^KO^* EAE mice had a small number of tdT^+^ cells, *Ifnar1^KO^* mice had a negligible number of tdT^+^ neurons and no tdT^+^ astrocytes, suggesting the direct stimulation of these populations with EAE is by type-I IFNs. Finally, we observed little-to-no CD4^+^ T cell infiltration in the brains of *Ifnar1^KO^*or *Ifngr1^KO^* mice (**Fig.S3A**).

**Figure 6.**
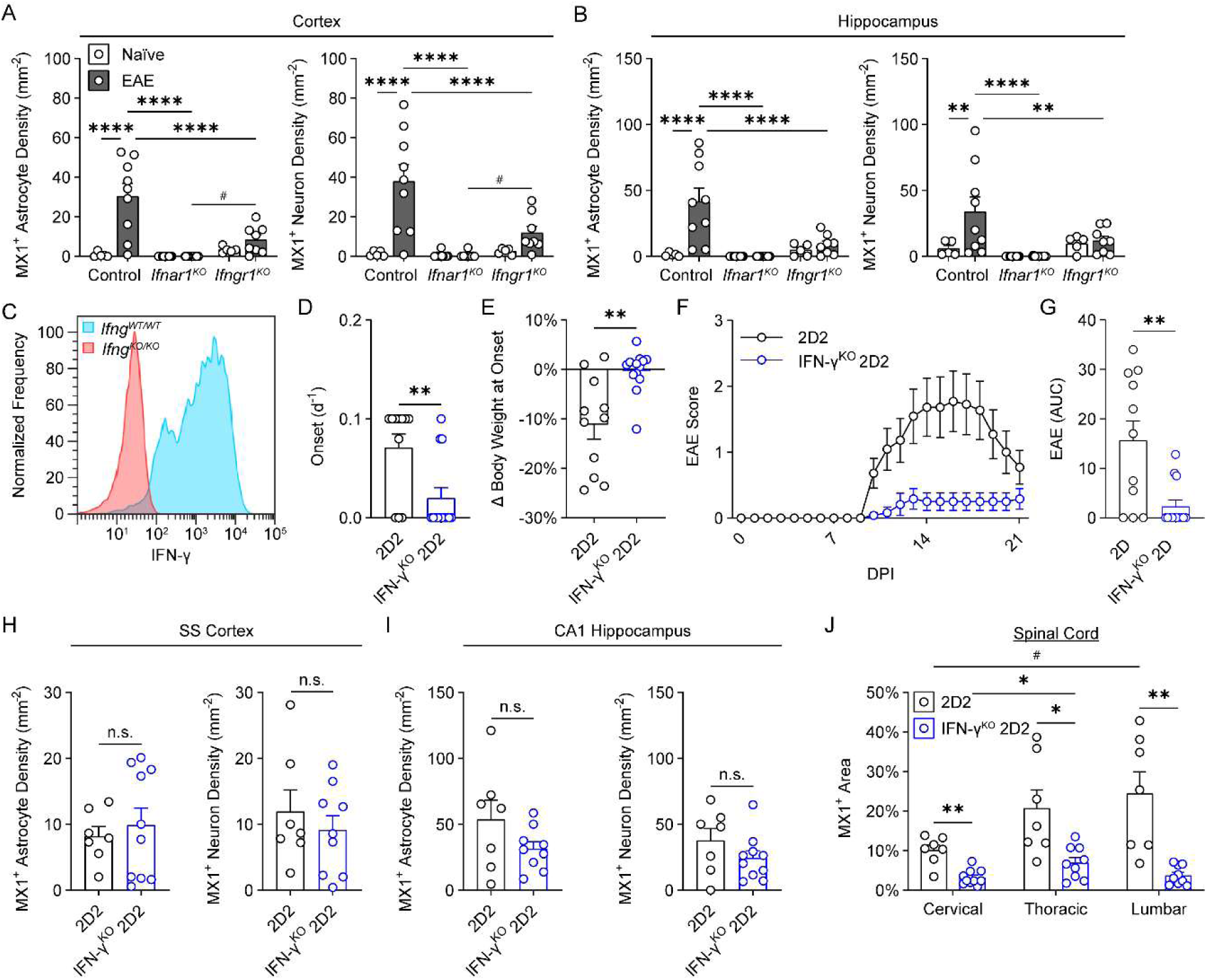
IFN-γ is necessary for EAE spinal cord pathology, but type-I IFN signaling is necessary for astrocyte and neuron pathology in the brain. (A&B) Quantification of MX1-tdT^+^ astrocyte and neuron density in the cortex and hippocampus of control, *Ifnar1^KO^*, and *Ifngr1^KO^* mice at 12-14 DPI. Genotype × Treatment Interaction for cortex astrocytes: F(2, 49) = 13.04, *P* < 0.0001; cortex neurons: F(2, 49) = 10.66, *P* = 0.0001, hippocampus astrocytes: F(2, 49) = 10.25, *P* = 0.0002; hippocampus neurons: F(2, 49) = 3.80, *P* = 0.0293. **(C)** Representative flow cytometry histogram confirming knockout of IFN-γ from Th1 2D2 cells. **(D)** Inverted day of EAE symptom onset (*i.e.*, no symptoms = 1/∞ = 0). **(E)** Change in body weight on day of symptom onset (or 12 DPI if no symptoms) compared to baseline. **(F)** EAE symptom scores measured each day. Treatment × Time Interaction: F(2, 37) = 8.88, *P* = 0.0012. **(G)** Area under the curve (AUC) for F. **(H&I)** Quantification of MX1-tdT^+^ astrocyte and neuron density in the SS cortex and CA1 region of the hippocampus at 21 DPI. **(J)** Quantification of MX1-tdT expression in the spinal cord at 21 DPI. Treatment × Region Interaction: F(1, 15) = 7.43, *P* = 0.0136. Data shown are mean ± SEM. \**P* < 0.05; \*\**P* < 0.01; \*\*\*\**P* < 0.0001; ^#^*P* < 0.10.

To isolate the role of IFN-γ production by 2D2 T_h_1 cells, passive EAE was induced using 2D2*^ΔIfng^* T_h_1 cells (**Fig. 6C**). While the majority of mice receiving normal 2D2 T_h_1 cells began to exhibit EAE symptoms at 10 DPI, most mice receiving 2D2*^ΔIfng^* T_h_1 cells did not exhibit any EAE symptoms (**Fig. 6D**). Additionally, while the 2D2 group lost body weight by the onset of symptoms, the 2D2*^ΔIfng^* group had not lost body weight by this time (**Fig. 6E**). Moreover, mice that had received 2D2*^ΔIfng^*T_h_1 cells that did develop EAE symptoms did not show paralysis past minor limpness of the tail (**Fig. 6F&G**). Interestingly, though 2D2*^ΔIfng^*mice did not develop clinical EAE symptoms, there was no difference in MX1-tdT expression in the brain between these groups (**Fig. 6H&I**). Contrarily, MX1-tdT expression in the spinal cord was attenuated by the knock-out of IFN-γ in the 2D2 T_h_1 used to induce EAE (**Fig. 6J**), and this IFN-γ^KO^ attenuated the infiltration of CD4^+^ T cells into the brain (**Fig.S3B&C**). These findings not only explain the lack of motor symptoms in these mice, but they also highlight a disconnect between the brain and spinal cord pathology of EAE.

### Astrocytes and microglia interact with CD4^+^ T cells at the medial edge of the fimbria at the onset of EAE symptoms

In addition to oligodendrocytes, ependymal cells, and vascular cells (*e.g.*, endothelial and mural cells), we identified three distinct clusters of astrocytes and two clusters of microglia within the fimbria in our spatial transcriptomics data (**Fig. 7A**). Importantly, the “Astrocyte 2” and “Microglia 2” clusters were unique (38% to 6% of astrocytes and 36% to 3% of microglia, respectively) to the EAE condition (**Fig. 7B**). Moreover, these “Microglia 2” cells were concentrated at the medial borders of the fimbria in EAE mice (**Fig. 7C**). Using immunofluorescence, we confirmed the concentration of MHC-II^+^ microglia/macrophages at this location (**Fig. 7D**). Compared to the other microglia of the fimbria, these “Microglia 2” cells expressed several inflammation-related transcripts and upregulated processes including complement-mediated synapse pruning (GO:0150062), complement activation, classical (GO:0006958) and alternative (GO:0030451) pathways, positive regulation of phagocytosis (GO:0050766), microglial cell activation involved in immune response (GO:0002282), positive regulation of neuroinflammatory response (GO:0150078), positive regulation of macrophage cytokine production (GO:0060907), cellular response to type-II IFN (GO:0071346), antigen processing and presentation of exogenous peptide antigen via MHC-II (GO:0019886), regulation of lymphocyte migration (GO:2000401), and positive regulation of T cell activation (GO:0050870), identifying these cells as antigen-presenting, IFN-γ-activated, neuroinflammatory microglia/macrophages (**Fig.7E&F**). Accordingly, the “Astrocyte 2” population upregulated the regulation of macrophage activation (GO:0043032) and immune response (GO:0006955) while downregulating the regulation of blood vessel remodeling (GO:0060312), cell junction organization (GO:0034330), L-glutamate import across plasma membrane (GO:0098712), and regulation of amyloid-β clearance (GO:1900221), reflecting inflammatory signaling with microglia/macrophages, dysfunction of the blood-brain barrier, decreased re-uptake of glutamate, and the accumulation of amyloid-β (**Fig.7G&H**). Spatial expression also confirms the concentration of relevant transcripts along this medial edge of the fimbria (**Fig.7I**).

**Figure 7.**
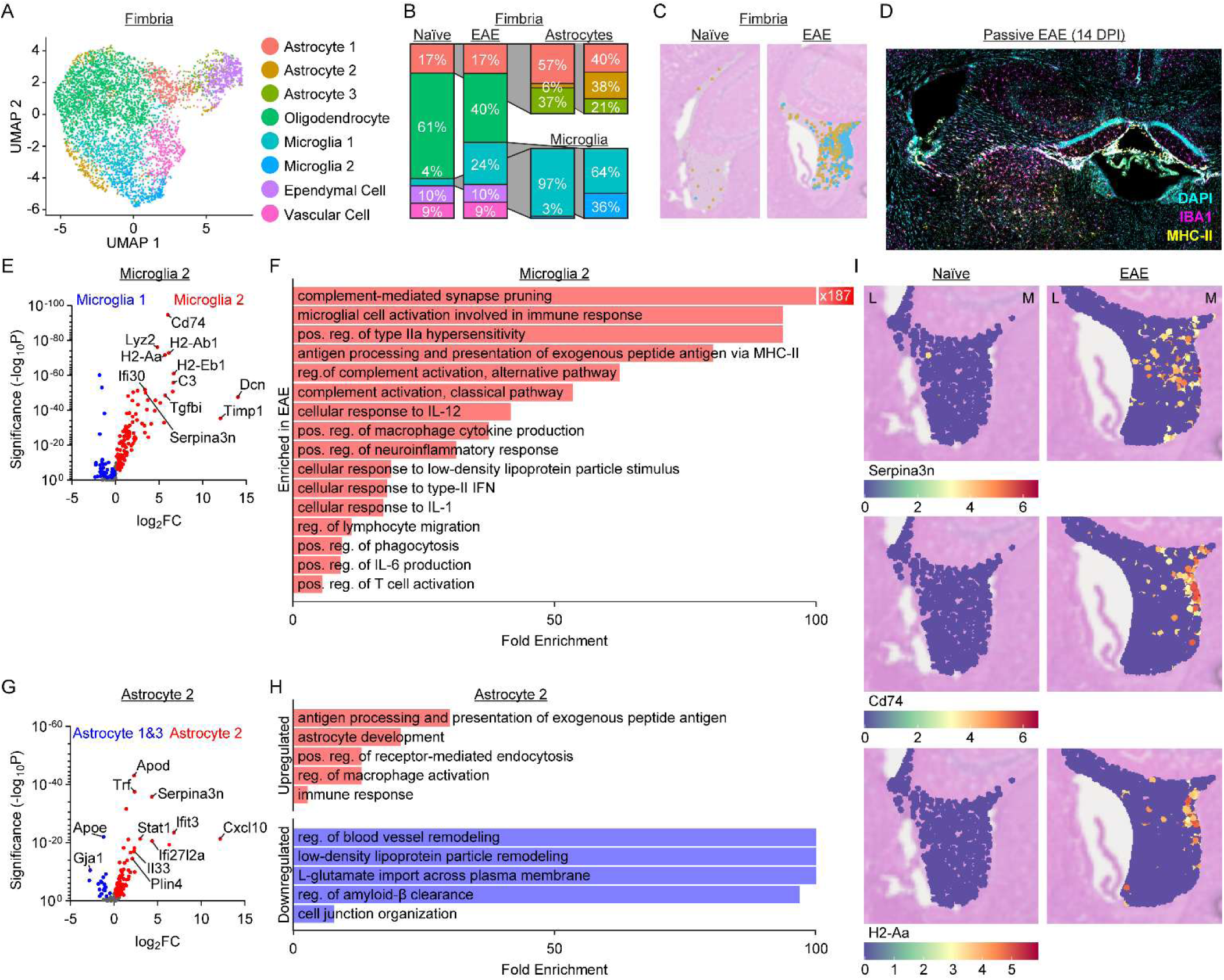
Astrocytes and microglia interact with CD4^+^ T cells at the medial edge of the fimbria at the onset of EAE symptoms. **(A)** Cell types and clustering of cells within the fimbria shown by UMAP plot. **(B)** Relative distribution of cell types and sub-clusters of astrocytes and microglia within the fimbria. **(C)** Representative spatial plots of “Astrocyte 2” and “Microglia 2” populations within the fimbria. **(D)** Representative image of microglia/macrophages at the medial borders of the fimbria expressing MHC-II. **(E)** Volcano plot of genes differentially expressed by “Microglia 2” cells compared to “Microglia 1” cells. **(F)** Significant enriched GO Biological Processes for “Microglia 2” DEGs. **(G)** Volcano plot of genes differentially expressed by “Astrocyte 2” cells compared to other astrocytes. **(H)** Significant enriched GO Biological Processes for “Astrocyte 2” DEGs. **(I)** Spatial expression of *Serpina3n*, *Cd74*, and *H2-Aa* within the fimbria.

## Discussion

While the major focus of MS and EAE research is on the spinal cord, studies have shown subtle pathology within the brain early in disease (Centonze et al., 2009; Politi et al., 2007). Here, we have shown autoreactive T cells activate IFN-dependent pathways in excitatory neurons within the SS cortex and CA1 region of the hippocampus early in EAE. Moreover, inhibition of type-I or -II IFN signaling attenuates the EAE-induced influx of T cells into the brain. Our data show microglia and astrocytes at the border between the myelin-rich fimbria and leukocyte-accessible third ventricle express elevated levels of MHC-II transcripts in EAE. The primary driver of this transcriptional change is IFN-γ stimulation (Steimle et al., 1994). Given our adoptively transferred 2D2 T_h_1 cells colocalize with myelin antigen-presenting cells and, as a result, produce IFN-γ, we interpret these data to show antigen recognition and IFN-γ signaling to microglia and/or astrocytes at the white matter-adjacent borders of the brain induces the infiltration of CD4^+^ T cells into the brain parenchyma.

Our sequencing data indicates that *Arc* expression is consistently elevated by EAE in both the SS cortex and CA1 hippocampus. While previous studies have shown a decrease in *Arc* expression with EAE in other regions of the brain (Centonze et al., 2009), dysregulation of *Arc* expression has been shown to be a potential causative factor in several neuropathologies (Shepherd and Bear, 2011). More importantly, *Arc* dysregulation is indicative of synaptic dysfunction, which correlates with the major dysregulated pathways we observed in our neuron sequencing. Additionally, we found that IFN-stimulated astrocytes in the fimbria downregulate transcriptional pathways associated with glutamate reuptake and cell junction organization, indicating a potential for increased excitotoxicity and dysfunctional calcium dynamics, respectively.

As with many autoimmune disorders, the prevalence of MS in women is 80% higher than in men (Jacobson et al., 1997). Sex-specific characteristics of the innate immune system, such as higher frequency of CD4^+^ T cells, greater phagocytic activity, and more efficient antigen presentation, may contribute to this bias (Klein and Flanagan, 2016). For example, our data show microglia in the fimbria activate an antigen-presenting, synapse-pruning transcriptional profile in response to EAE. Given microglia in the female brain are already skewed toward this phenotype, they may be more susceptible to this pathology.

Astrocytes have previously been shown to perform several functions, such as antigen presentation and phagocytosis, typically attributed to microglia in the brain (Fontana et al., 1984; Guo et al., 2026; Lee et al., 2021; Ponath et al., 2017; Stüve et al., 2002). While we did not directly observe phagocytic activity by astrocytes in this study, our identification of MHC-II expression by astrocytes in the brain is consistent with recent work performed by Lee *et al*. (2026). Additionally, astrocytic IFN-α overexpression has been shown to induce inflammation and neurodegeneration within the CNS (Akwa et al., 1998). Others have shown that inhibition of IFN-α improves cognitive function and attenuates loss of dendritic arborization in a glutamate-NMDA receptor-dependent manner in a mouse model of human immunodeficiency virus encephalitis (Sas et al., 2009). Apart from IFN signaling, astrocytes have been shown to modulate synapse engulfment and neuronal circuit development in an IL-33-dependent manner (Han et al., 2023; Nguyen et al., 2020; Vainchtein et al., 2018). Additionally, within the SS cortex specifically, Moreno-Garcia et al. found that EAE caused astrocyte calcium signals of increased duration and decreased amplitude, and these astrocytic alterations coincided with the onset of neurological symptomatology (2025). Taken together, these studies indicate astrocytic type-I IFN production causes dysfunction of neurons and astrocytes themselves at excitatory synapses. Moreover, our data suggest astrocyte calcium dysregulation and decreased glutamate uptake, neuron glutamate-NMDA receptor dysregulation, and microglial synaptic pruning all play a role in this synaptic pathology.

In summary, we have identified the activation of transcriptional pathways associated with antigen presentation by microglia and astrocytes within the fimbria as a key gateway for IFN-dependent leukocyte infiltration into the brain parenchyma in EAE and, potentially, MS. IFN signaling within the brain supports our transcriptional data suggesting excessive synaptic pruning by microglia, and this is likely caused by synaptic dysfunction and IFN-stimulation of excitatory neurons (Gillani et al., 2024). Thus, this study highlights key steps in CNS autoimmunity that lead to neuronal pathology and, likely, cognitive symptoms in patients with MS and provides several avenues for potential therapeutic intervention.

## Materials & Methods

### Transgenic Mouse Lines

Transgenic mice were purchased from The Jackson Laboratory and bred at Duke University facilities. 2D2 TCR mice (JAX#006912) were bred with (1) C57B/6J CD45.1 mice (JAX#002014) to produce 2D2^CD45.1/2^ mice expressing both CD45.1 and CD45.2, (2) UBC-GFP mice (JAX#004353) to produce 2D2^GFP^ mice expressing GFP in hematopoietic cells, or (3) *Ifng^KO^* mice (JAX#002287) to produce 2D2*^ΔIfng^*mice lacking IFN-γ expression in all cells. Mx-CRE mice (JAX#003556) were bred with Ai9 mice (JAX#007909) to produce MX1^tdT^ mice expressing fluorescent tdTomato protein in interferon-stimulated cells. These mice were then bred with *Ifnar1^KO^* mice (JAX#028288) or *Ifngr1^KO^*mice (JAX#003288) to produce MX1^tdT^ mice with either *Ifnar1* or *Ifngr1* knocked out on all cells, respectively. Both male and female mice were used for experiments, but no effect of biological sex was detected throughout the study. All mice were housed under barrier conditions with a 12/12 h light/dark cycle and *ad libitum* access to rodent chow and water. All procedures were performed in accordance with the National Institutes of Health Guide for the Care and Use of Laboratory Animals and were approved by Duke Institutional Animal Care and Use Committee.

### Induction of Active EAE

To induce active EAE, mice were immunized using subcutaneous injection of MOG_35-55_ peptide emulsified in Complete Freund’s adjuvant (Hooke Laboratories). On the day of immunization and 24 h later, mice received an intraperitoneal injection of 120 ng pertussis toxin (in PBS). Mice were observed daily and euthanized at appropriate time points. Symptoms were characterized as follows: 0.0 – tail is erect; 0.5 – tip of tail is limp; 1.0 – tail is limp; 1.5 – hind leg inhibition; 2.0 – hind legs are weak and toes of one paw are dragged while walking; 2.5 – hind legs are dragging; 3.0 – hind legs are paralyzed; 3.5 – unable to right self when placed on side or hind quarters have flattened appearance; 4.0 – front legs are partially paralyzed; 4.5 – mouse is not alert; 5.0 – mouse is moribund.

### Immunohistochemistry

For immunohistochemistry (IHC), mice were transcardially perfused with ice-cold PBS and 4% paraformaldehyde (PFA). Tissue was stored in 4% PFA overnight. Brains and spinal cords were dehydrated with 30% sucrose, frozen over dry ice, and sectioned coronally at 30 μm on a Leica CM1950 cryostat. Sections were stored in a solution of 30% ethylene glycol, 30% polyethylene glycol, and 40% 0.2 M phosphate buffer for IHC. Dura mater tissue was dissected from the skull cap and stored in PBS for IHC.

Tissue was washed with PBS and blocked/permeablized with 5% normal donkey serum, 0.5% Triton X-100, and 1% BSA in PBS for 1 h prior to incubation overnight at 4°C with antigen-specific primary antibodies (in blocking/permeablization solution). Tissue was then washed with PBS and incubated with fluorophore-conjugated secondary antibodies for 2 h. Tissue was then washed with PBS, stained with DAPI (if applicable), mounted onto slide glass, and coverslipped with Fluoromount-G (SouthernBiotech). Tile scan images were collected at 10X magnification on an EVOS FL Auto epifluorescence microscope (Thermo Fisher). Images were analyzed using ImageJ software (NIH). High-magnification images were acquired using an upright Leica SP8 confocal microscope with an oil-immersion 63X objective.

Primary antibodies used were 1:200 rat anti-CD4 (Invitrogen 14-0042-82), 1:200 rat anti-CD68 (abcam ab53444), 1:1,000 rabbit anti-IBA1 (Wako Chemicals 019-19741), 1:250 rat anti-MHC-II (Invitrogen 14-5321-82), 1:2,000 rabbit anti-NeuN (abcam ab177487), and 1:100 goat anti-SOX9 (BioLegend 110723).

### Cell Density Analysis

MX1^+^ cell density was quantified in the cortex and hippocampus by region. Cell counts were divided by the measured tissue area in coronal sections. Neurons were identified by NeuN expression, astrocytes by SOX9 expression, CD4^+^ T cells by CD4, and microglia/macrophages by IBA1.

### T_h_ Cell Culture & Passive EAE

To induce passive EAE, naïve CD4^+^ T cells were isolated from the lymph nodes and spleen of 2D2 TCR mice using the MojoSort™ Mouse CD4 Naïve T Cell Isolation Kit (BioLegend). T cells were incubated at 37°C and 5% CO_2_ in RPMI medium supplemented with 10% FBS, 0.1 μg/mL anti-CD3, 0.1 μg/mL anti-CD28, 2 μg/mL anti-IL-4, 50 U/mL IL-2, and 10 ng/mL IL-12 for Th1 skewing conditions. After five days, 5×10^6^ cells were injected intravenously (i.v.) via tail vein into each mouse. A sample of cells was used to confirm Th1 activation (expression of CD4, IFN-γ, and T-bet) via flow cytometry after PMA/ionomycin stimulation. Mice were observed daily and euthanized at appropriate time points. Symptoms were characterized as detailed above.

### Tissue Digestion & Flow Cytometry

For flow cytometry, the dura mater was isolated from the skull cap and digested with collagenase D and DNase prior to homogenization over a 70-μm tissue screen. Lymph nodes and spleens were homogenized over 70-μm tissue screens, and spleens were further processed with ACK Lysing Buffer (Quality Biological) to remove red blood cells. Finally, blood was processed with ACK Lysing Buffer. Cells were labeled with fluorophore-conjugated, antigen-specific antibodies and resuspended in FACS buffer (2% FBS in PBS) for analysis using a FACSLyric flow cytometer (Becton Dickinson). “Single-stain” and “fluorescence minus one” controls were run for proper quantification of fluorescence. Data were analyzed using FlowJo software (Becton Dickinson).

### Spatial RNA Sequencing & Analysis

Spatial RNA sequencing was performed on 10-µm coronal brain sections from mice at the onset of passive EAE symptoms. Brains were processed as for IHC prior to mounting onto Visium HD specialized slides with CytAssist (10X Genomics). Hematoxylin and Eosin (H&E) staining was performed and reference images obtained prior to Visium HD probe capture and sequencing. Sequencing data were processed using Space Ranger (10X Genomics) and ENACT cell segmentation software (Kamel et al., 2025). Identification of MX1 reporter-rich regions of interest was made using IHC on sequential coronal sections.

Sequencing data was processed and analyzed using the Seurat Bioconductor package in R (Hao et al., 2024). Differentially expressed genes (DEGs) were determined by (1) expression in at least 10% of either population, (2) Benjamini-Hochberg adjusted *P*-value less than 0.05, and (3) an absolute log_2_(fold change) greater than 0.1. Statistical overrepresentation of Gene Ontology (GO) Biological Processes for differentially expressed genes was calculated using the PANTHER (<u>P</u>rotein <u>AN</u>alysis <u>TH</u>rough <u>E</u>volutionary <u>R</u>elationships) Classification System (Thomas et al., 2022). All Biological Processes referenced have adjusted *P*-values < 0.05.

### Statistical Analysis

Single-variable, two-level analyses were conducted using a two-tailed Welch-corrected *t*-test. Single-variable, analyses with more than two levels were conducted using a one-way <u>an</u>alysis <u>o</u>f <u>va</u>riance (ANOVA) followed by Šídák’s multiple comparisons test. Multi-variate analyses were conducted using two-way ANOVA followed by Šídák’s multiple comparisons test (if applicable). RNA-sequencing differential expression testing was performed using the non-parametric Wilcoxon rank sum test.

## Data Availability

Bioinformatics data from spatial RNA-sequencing in this study are available via the Gene Expression Omnibus (GSE346572). Other data may be made available from the corresponding author upon reasonable request. All other data underlying the findings of this study can be made available upon request to the corresponding author.

## Acknowledgements

This work was supported by the National Institutes of Health R01NS123084. We would like to acknowledge the assistance of the Molecular Genomics Core at the Duke Molecular Physiology Institute, Duke University School of Medicine, for the generation of Visium HD RNA sequencing data, which we then processed using an allocation from the Duke Compute Cluster.

Disclosures: A.J. Filiano had intellectual property licensed to Cryo-Cell International. The agreement ended in 2026.

## Abbreviations

CNS: central nervous system
DEG: differentially expressed gene
DPI: days post immunization
EAE: experimental autoimmune encephalomyelitis
IHC: immunohistochemistry
MOG: myelin oligodendrocyte glycoprotein
MS: multiple sclerosis
PFA: paraformaldehyde
PTX: pertussis toxin
SS: somatosensory
tdT: tdTomato
T_h_1: T helper 1

---

**Figure S1.**
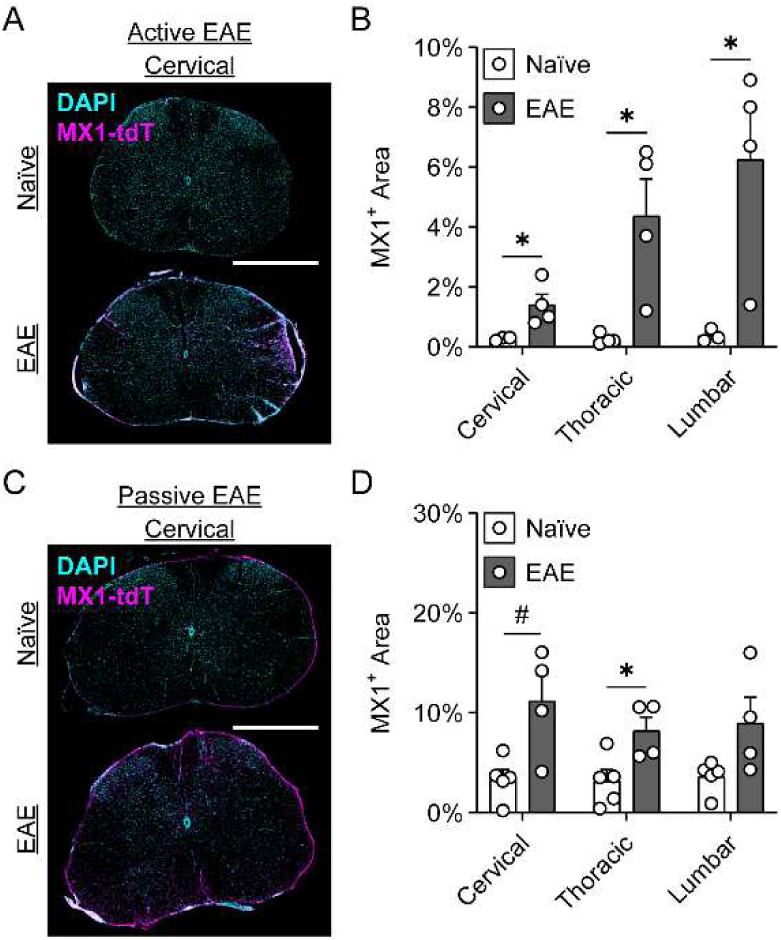
IFN signaling increases in the spinal cord at the onset of EAE symptoms. **(A)** Representative spinal cord sections from active EAE mice at 14 DPI. Scale bar = 1 mm. **(B)** Quantification of MX1-tdT expression in the spinal cord of active EAE mice. Main Effect of Treatment: F(1, 6) = 18.01, *P* = 0.0054. **(C)** Representative spinal cord sections from passive EAE mice at 12 DPI. Scale bar = 1 mm. **(D)** Quantification of MX1-tdT expression in the spinal cord of passive EAE mice. Data shown are mean ± SEM. \**P* < 0.05; ^#^*P* < 0.10.

**Figure S2.**
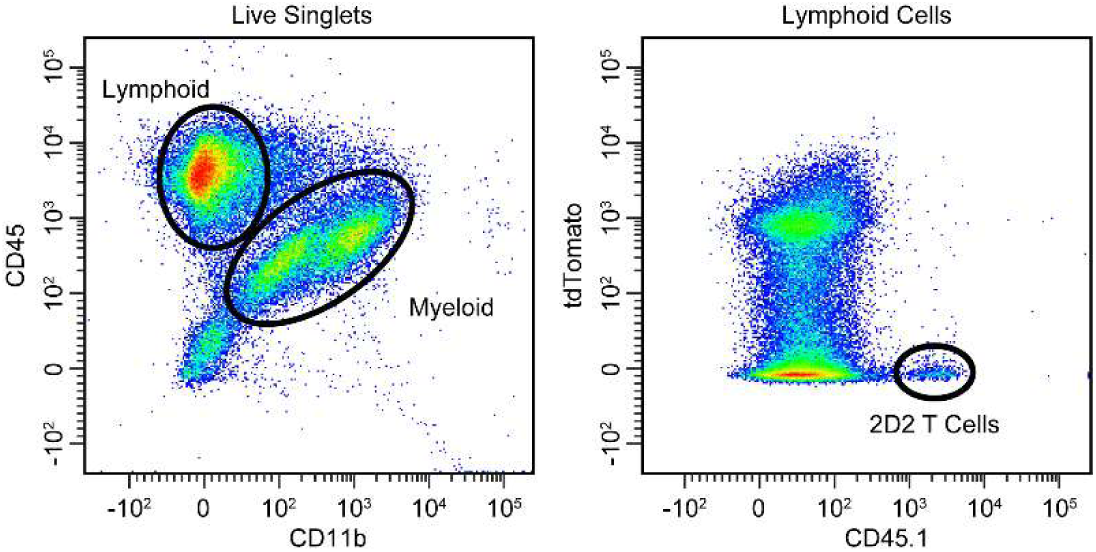
Representative flow cytometry gating strategy to identify MX1-tdT^+^ 2D2 T cells. Lymphoid cells were defined as CD45^+^/CD11b^-^ populations. 2D2 cells were defined as lymphoid cells that were tdT^-^/CD45.1^+^.

**Figure S3.**
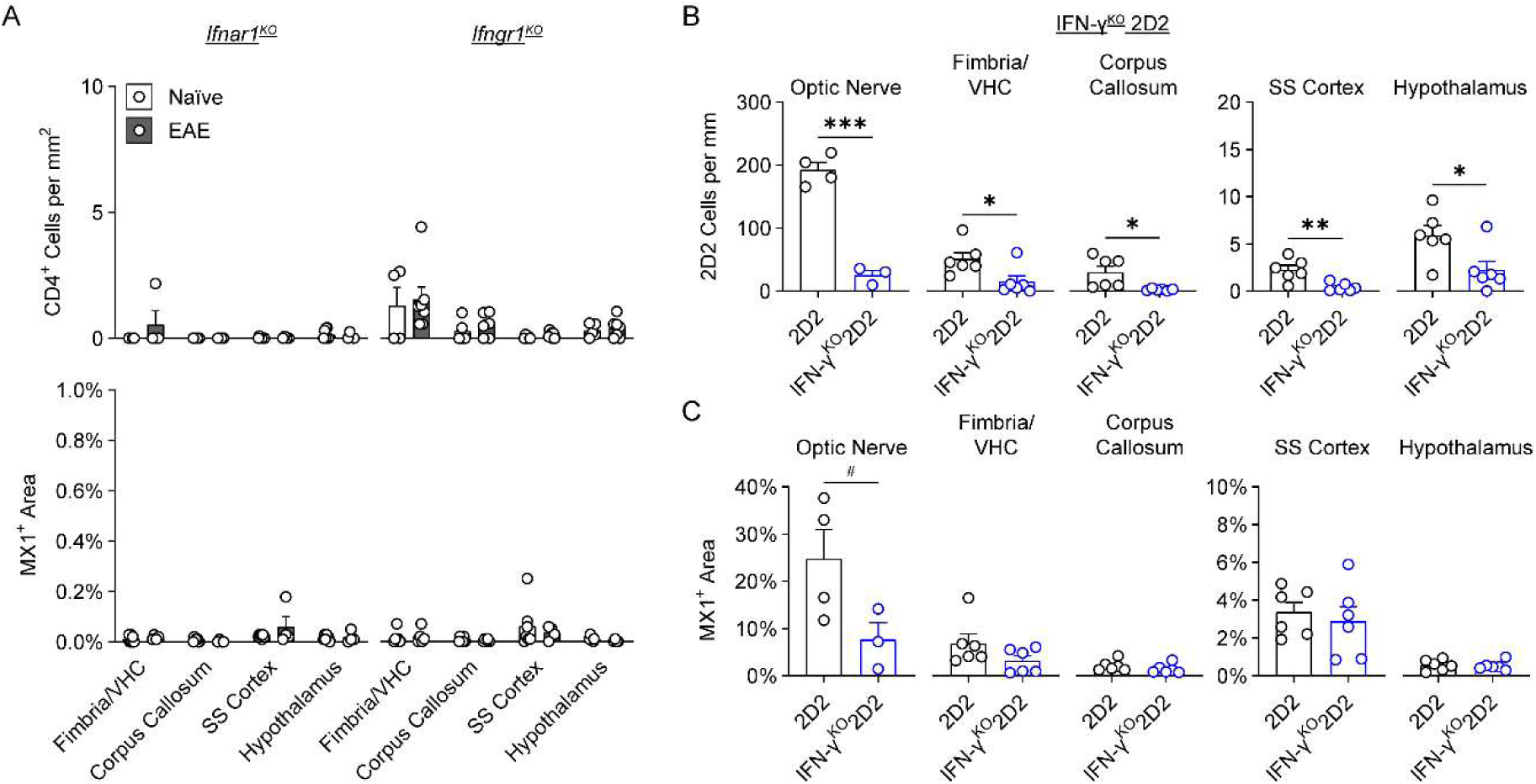
Attenuation of IFN-signaling in EAE prevents the infiltration of CD4^+^ T cells into the brain. **(A)** Quantification of CD4^+^ T cell infiltration and MX1-tdT expression in various white matter and gray matter regions of *Ifnar1^KO^* and *Ifngr1^KO^* mice with and without EAE induction. **(B&C)** Quantification of CD4^+^ T cell infiltration and MX1-tdT expression in various white matter and gray matter regions of mice that received 2D2 or IFN-γ^KO^ 2D2 Th1 cells. Data shown are mean ± SEM. \**P* < 0.05; \*\**P* < 0.01; \*\*\**P* < 0.001.

